# Inhibitory effect of capsule on natural transformation of *Streptococcus pneumoniae*

**DOI:** 10.1101/2025.05.07.652784

**Authors:** Sheya Xiao Ma, Hannes Eichner, Michael Cammer, Jeffrey N. Weiser

## Abstract

The capsule of *Streptococcus pneumoniae* (*Spn*) is highly heterogeneous based on expression of distinct polysaccharides. *Spn* transformation, controlled by the Com regulon, has been predominantly focused on unencapsulated laboratory strains. However, genomic studies revealed different rates of recombination events in clinical isolates of different serotypes. As these isolates were genetically distinct beyond capsule-encoding genes, the exact relationship between transformation and capsule remains unclear. Herein, we compared the transformability of a collection of isogenic capsule-switch strains. Strains with different capsule types and amounts significantly differed in their transformation frequency, with the unencapsulated strain having a higher frequency compared to encapsulated strains. A GFP-reporter of each strain monitoring the expression of a Com regulon-controlled gene showed similar kinetics, indicating differences in transformability were due to processes downstream of competence activation. The Com pilus, induced by competence, binds and takes in the donor DNA, and is the central component of the transformation apparatus. The surface exposure of Com pilus significantly differed among serotypes with highly transformable strains having more cells binding ComGC antibody. Further, electron microscopy demonstrated that transformability correlated with the proportion of cells bearing a Com pilus, which was affected by both the presence of capsule and serotype. Additionally, the unencapsulated strain displayed longer pili than encapsulated strains. Examination of capsule porosity revealed that serotypes with higher transformation frequencies had more porous capsules. Together, these results indicate that the capsule interferes with the assembly of Com pilus, thereby inhibiting the natural transformation of *Spn*.

**IMPORTANCE:** The capsule is a major virulence factor of *Streptococcus pneumoniae* (*Spn*), providing a physical shield and exhibiting extensive diversity across at least 100 serotypes. Although natural transformation of *Spn* has predominantly been characterized in unencapsulated laboratory strains, clinical encapsulated isolates also exhibit transformability and demonstrate varied recombination rates during host carriage. We utilized otherwise genetically identical capsule-switch strains to isolate the effect of capsule on transformation. We demonstrate serotype- and quantity-dependent inhibition of transformation by the capsule, mediated through hindrance with the transformation pilus assembly and function. This study challenges the paradigm that unencapsulated laboratory strains fully recapitulate natural transformation dynamics. By redefining capsule as a multifunctional modulator of *Spn* biology, balancing virulence and adaptability, our findings advance our understanding of pneumococcal evolution.

## INTRODUCTION

The major virulence factor of *Streptococcus pneumoniae* (*Spn*, the pneumococcus) is its capsule, a polysaccharide coat of varying thickness, which impedes mucus entrapment and shields the bacterium from opsonophagocytic killing [1]. Over 100 structurally and antigenically distinct capsule types, or serotypes, determined by heterogeneity in the capsular polysaccharide synthesis (*cps*) locus, have been described [2–4]. Capsular polysaccharides are the basis of all current pneumococcal conjugate vaccines (PCVs), which are highly effective against included serotypes [5]. However, due to serotype diversity, the limited serotype coverage (≤21 types) of PCVs exerts selective immune pressure favoring nonvaccine serotypes [6, 7]. Additionally, “capsule switch” vaccine escape variants, may emerge by genetic exchange involving the *cps* genes between nonvaccine and vaccine serotypes [8–13]. This phenomenon that threatens to erode vaccine efficacy, highlights the clinical consequences of the genetic plasticity of *Spn*, which is attributed to its natural transformability.

Since Griffith’s seminal discovery in 1928, *Spn* has served as a model organism for natural genetic transformation [14]. The state of competence for DNA uptake is a transient, tightly regulated process governed by a competence regulon (Com regulon) activated via quorum sensing [15]. The competence-stimulating peptide (CSP) encoded by *comC* is secreted via the ABC-transporter, ComAB, and sensed by the two-component regulatory system, ComDE. Binding of CSP to the membrane-spanning histidine kinase, ComD, triggers autophosphorylation of ComD and subsequent phosphorylation of its cognate response regulator, ComE. Activated ComE induces early competence genes, including *comX*, which in turn activates late competence genes responsible for DNA uptake, homologous recombination, and fratricide. A key component of the DNA uptake machinery is the competence pilus (Com pilus, transformation pilus), which is formed exclusively in competent cells [16]. These dynamic filamentous appendages, which extend and retract actively on the cell surface, bind extracellular DNA and facilitate its transport into the cell [16–18]. Captured double-stranded DNA is transferred to the DNA receptor ComEA, where one strand is degraded by the EndA endonuclease [19, 20]. The resulting single-stranded DNA traverses the ComEC pore into the cytosol allowing for its integration into the genome via homologous recombination [15]. Genetic recombination through natural transformation is a pivotal evolutionary mechanism enabling *Spn* to rapidly adapt to environmental pressures, including to antibiotics [21]. Notably, *Spn* ranks among the leading pathogens associated with antimicrobial resistance-related deaths, imposing a substantial global healthcare burden [22].

Historically, the pioneering studies from the lab of Oswald Avery bifurcated research on *Spn* into studies focusing either on the capsule and its role in virulence or the molecular mechanisms of transformation [23]. For decades, these fields have evolved separately, with transformation studies predominantly utilizing unencapsulated mutants selected for their robust transformability *in vitro*. However, encapsulated *Spn* strains, which are physiologically relevant in clinical contexts, are also naturally transformable and have contributed to the emergence of vaccine escape variants and multidrug resistance. While it has long been recognized that transformation is far more challenging to demonstrate in the laboratory setting with densely encapsulated strains compared to less encapsulated or unencapsulated strains, the underlying mechanisms have not been described [24, 25]. Comparative genomic studies have reported that clinical *Spn* isolates clustered by different serotypes exhibit a highly variable frequency of recombination events [26]. This finding suggests potential effects of serotype on genetic recombination through transformation that impact the evolution of different *Spn* lineages. However, considering the marked differences in serotype prevalence (and, thus, opportunities for genetic exchange) and the differing genetic backgrounds of clinical isolates, the exact relationship between capsule and transformation remains unclear.

Here, we systematically investigate the role of capsule in *Spn* transformation using a panel of isogenic capsule-switch mutants. We compare the transformation frequency among these strains and elucidate the mechanisms by which capsule and its type affect natural transformation, bridging the gap between virulence characteristics and genetic adaptability in *Spn* biology.

## RESULTS

### Different transformation frequencies among capsule-switch strains

To evaluate capsule-specific effects on pneumococcal transformation, we constructed a panel of isogenic mutants derived from a T4*^cps^*^-^ genetic background, expressing distinct capsule types or varying capsule amounts (Table 1) [27]. The panel included commonly circulated serotypes that are covered by PCVs (type 4, 6A, 7F, 14, 19F, 23F), serotypes with neutral surface charges (type 7F and 14), and those with extremely high or low recombination rates in genomic studies (type 16F and 35B) [26].

**Table 1.**
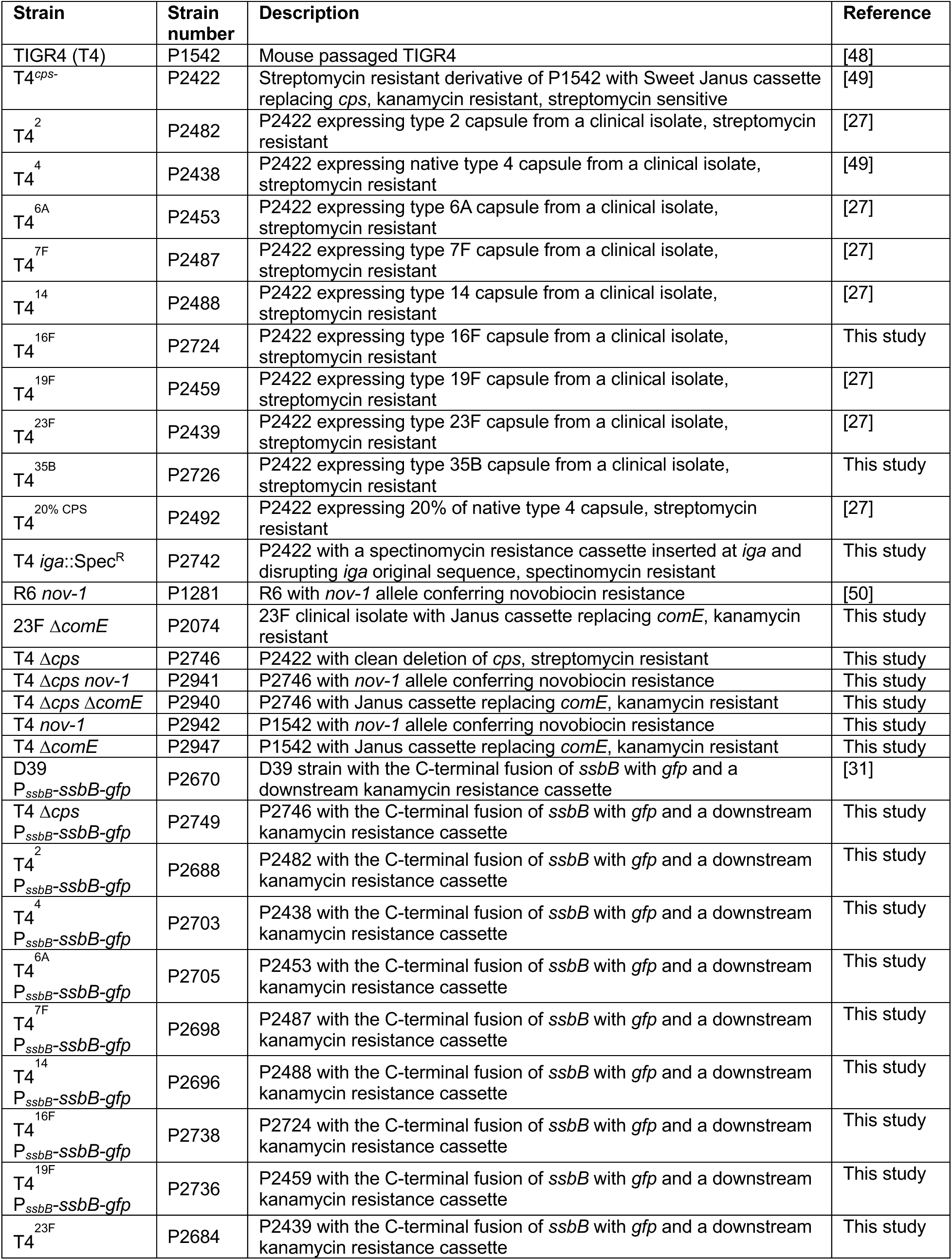
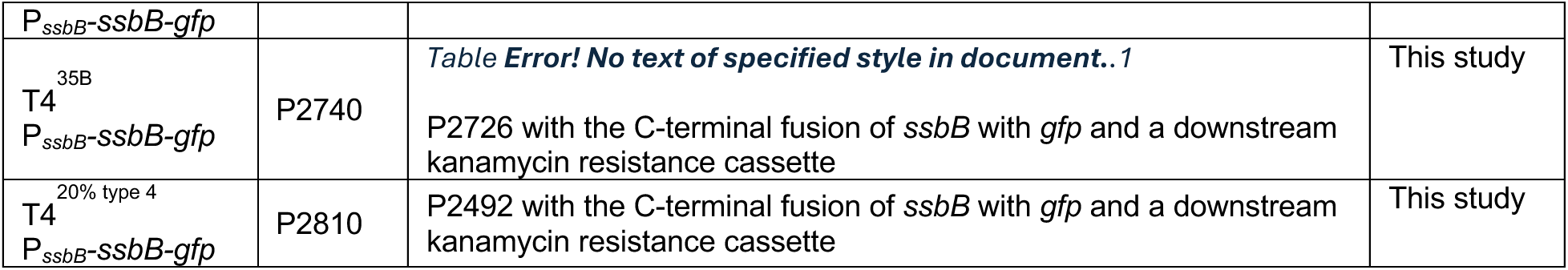
Bacterial strains used in this experimental work.

In pilot studies, we compared the concentration of CSP required to induce competence. At 150 ng/mL, significant differences in transformation frequencies among strains emerged, without major alterations upon further CSP concentration increases (Fig. 1a). Therefore, 150 ng/mL CSP was selected for subsequent analyses to induce competence.

**Fig 1.**
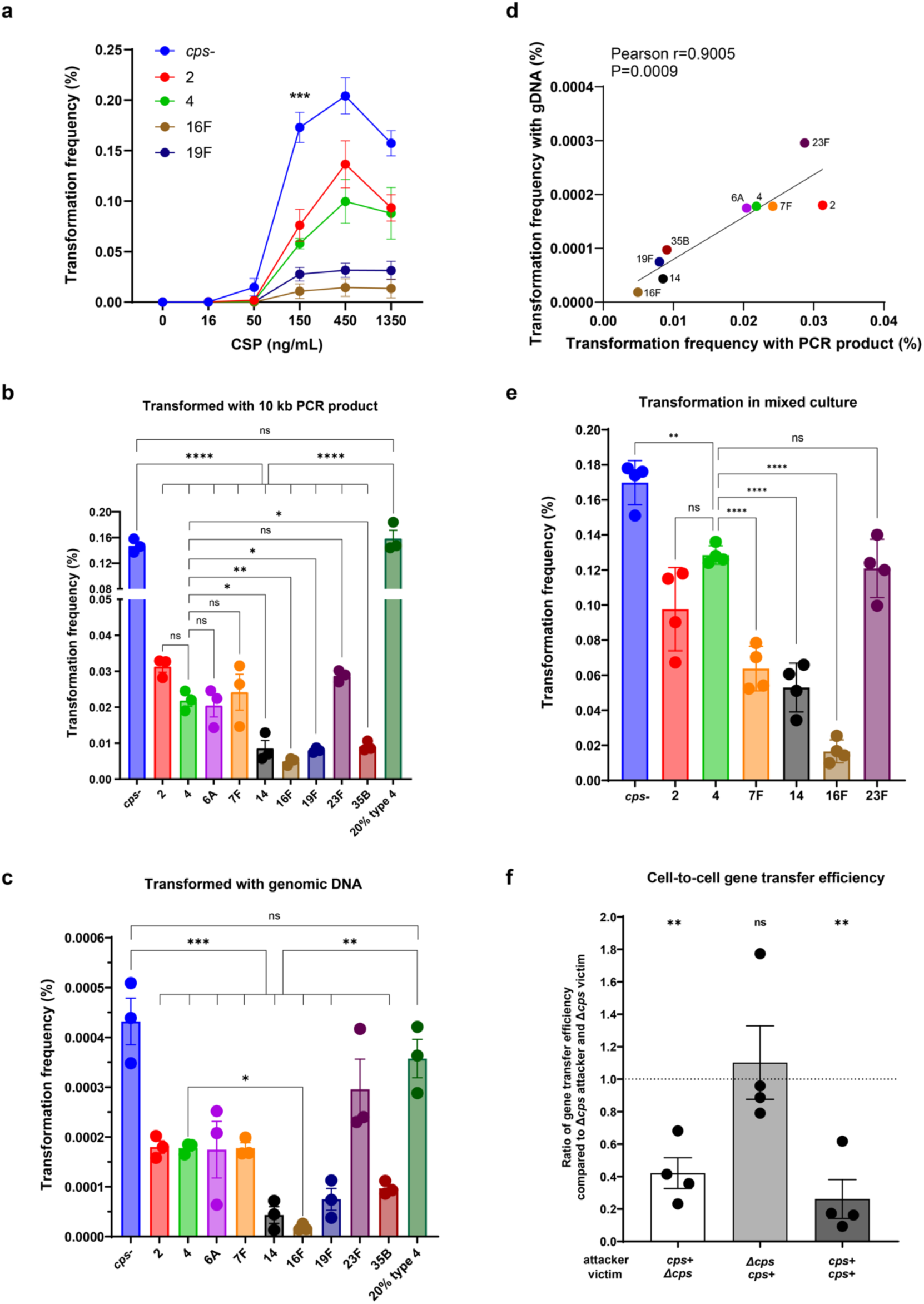
Transformation frequency of isogenic strains with different types or amounts of capsule. (a) Transformation frequency of capsule-switch strains responding to different CSP dosage. The transformation frequency with 150 ng/mL CSP was compared between strains using ordinary one-way ANOVA with Tukey’s multiple comparisons test. For brevity, only significance compared to *cps*- was shown on the graph. (b-c) Transformation frequency of capsule-switch strains with 150 ng/mL CSP and a Spec^R^ PCR product (b) or genomic DNA bearing the same selection marker (c). Transformation frequency of the unencapsulated strain (*cps*-) or the strain with 20% type 4 capsule was compared to the rest strains using the ordinary one-way ANOVA with Šídák’s multiple comparisons test. Transformation frequency of strains with different capsule types were compared using the ordinary one-way ANOVA with Tukey’s multiple comparisons test. For brevity, only comparisons to *cps*-, type 4, or 20% type 4 were shown on the graph. (d) Correlation between transformation frequency of strains bearing different capsule types transformed with PCR product and transformed with genomic DNA. Pearson correlation coefficients were computed. (e) Transformation frequency of capsule-switch strains with 150 ng/mL CSP and Spec^R^ PCR product in mixed culture. Culture of individual strain were mixed to perform transformation simultaneously. Colonies of each serotype were distinguished by colony immunoblot. Statistical significance was determined using an ordinary one-way ANOVA with Tukey’s multiple comparisons test. For brevity, only comparisons to type 4 were shown on the graph. (f) Efficiency of cell-to-cell gene transfer with Δ*cps* or *cps*+ attacker and victim. After incubating Nov^R^ attacker with Kan^R^ victim, the proportion of resulting dual-resistant colonies was calculated. Data show the ratio of dual-resistant proportion of each group compared to the control group, Δ*cps* attacker and Δ*cps* victim. The ratio was compared to a hypothetical mean of 1 using the one sample t test. Strains are designated by their capsule type or amount for brevity. Data show mean ± SEM from at least 3 independent experiments. ns, not significant; *, P≤0.05; **, P≤0.01; ***, P≤0.001; ****, P≤0.0001.

Next, we compared the transformation frequency of the entire panel of capsule-switch strains transformed with either 10 kb PCR product or whole genomic DNA containing the same selection marker. As expected, the *cps*-strain displayed much higher transformation frequency compared to fully encapsulated strains, whether using PCR product or genomic DNA (Fig. 1b and 1c). A *cps* promoter construct with ∼20% of WT levels of type 4 capsule resembled the *cps*-strain in transformation frequency (Fig. 1b and 1c), suggesting a threshold effect of capsule amount on transformation inhibition. Additionally, serotypes 2, 4, 6A, 7F, and 23F consistently exhibited higher transformation frequencies compared to serotypes 14, 16F, 19F, and 35B, with up to a 10-fold difference (Fig. 1b and 1c). There was a proportional relationship between transformation efficiencies with PCR product or genomic DNA (Fig. 1d).

To further confirm serotype-specific effects on transformation, we examined the transformation frequency of each strain under identical transformation conditions. Cultures of individual strains at the same density were equally mixed prior to transformation and colonies of each serotype were distinguished by colony immunoblot using serotype-specific antisera. The *cps*-strain maintained the highest transformation frequency, and consistent serotype-specific differences persisted in mixed conditions, with up to a 13-fold variation between encapsulated strains (Fig. 1e). Thus, these results validated that the difference in transformation frequency was serotype-specific, independent of other cellular or transformation condition variables.

Since, under natural conditions, *Spn* likely acquires DNA from sibling cells via intra-strain competition, we assessed capsule effects on cell-to-cell gene transfer in a modified fratricide assay [28–30]. We constructed a panel of competence-proficient, ‘attacker’ strains (novobiocin-resistant), and competence-deficient, ‘victim’ strains (kanamycin-resistant) in the presence or absence of a native capsule (Table 1). After co-incubation of attacker and victim in the presence of CSP, the gene transfer efficiency from victim to attacker was assessed by dual resistance emergence. Compared to the control group where both attacker and victim were unencapsulated, there was a significant reduction of gene transfer efficiency when the attacker was encapsulated, while victim encapsulation alone did not impair gene transfer rates (Fig. 1f). These results indicate capsule interferes with DNA internalization by recipients rather than DNA release from donors during intercellular gene transfer. This was further validated by the fact that dual encapsulation mirrored attacker-only encapsulation outcomes, but with lower efficiency than the victim-only encapsulation outcomes (Fig. 1f). Collectively, our findings establish that capsule suppresses natural transformation in a serotype- and quantity-dependent manner, predominantly by hindering recipient cell DNA internalization during horizontal gene transfer.

### Competence induction cannot explain capsule-dependent transformation differences

Next, we examined whether the capsule affects competence induction, hypothesizing that it might shield against CSP uptake. The kinetics of competence development was monitored using a translational *ssbB-gfp* fusion reporter in each capsule-switch strain using flow cytometry (Table 1) [31]. The late competence gene *ssbB* serves as a reliable indicator of competence induction due to its high expression and strong correlation with transformation [32–34]. All capsule-switch strains displayed similar competence kinetics, with the proportion of competent cells peaking between 20–30 min post-induction (Fig. 2a). Notably, despite a minor discrepancy of type 35B, these high competence levels starkly contrasted with the variable transformation frequencies (Fig. 1b and 2b). No correlation was found between either the proportion of competent cells or mean fluorescence intensity of the reporter (Fig. 2d and 2e) with transformation frequency (Fig. 2c and 2f), suggesting that the level of competence does not account for differences in transformation frequency in strains with differing types or amounts of capsule. In addition, we excluded fitness defects as a potential factor, since all strains exhibited similar growth characteristics (Fig. S1a) and there was no correlation between maximum growth rate and transformation efficiency (Fig. S1b).

**Fig 2.**
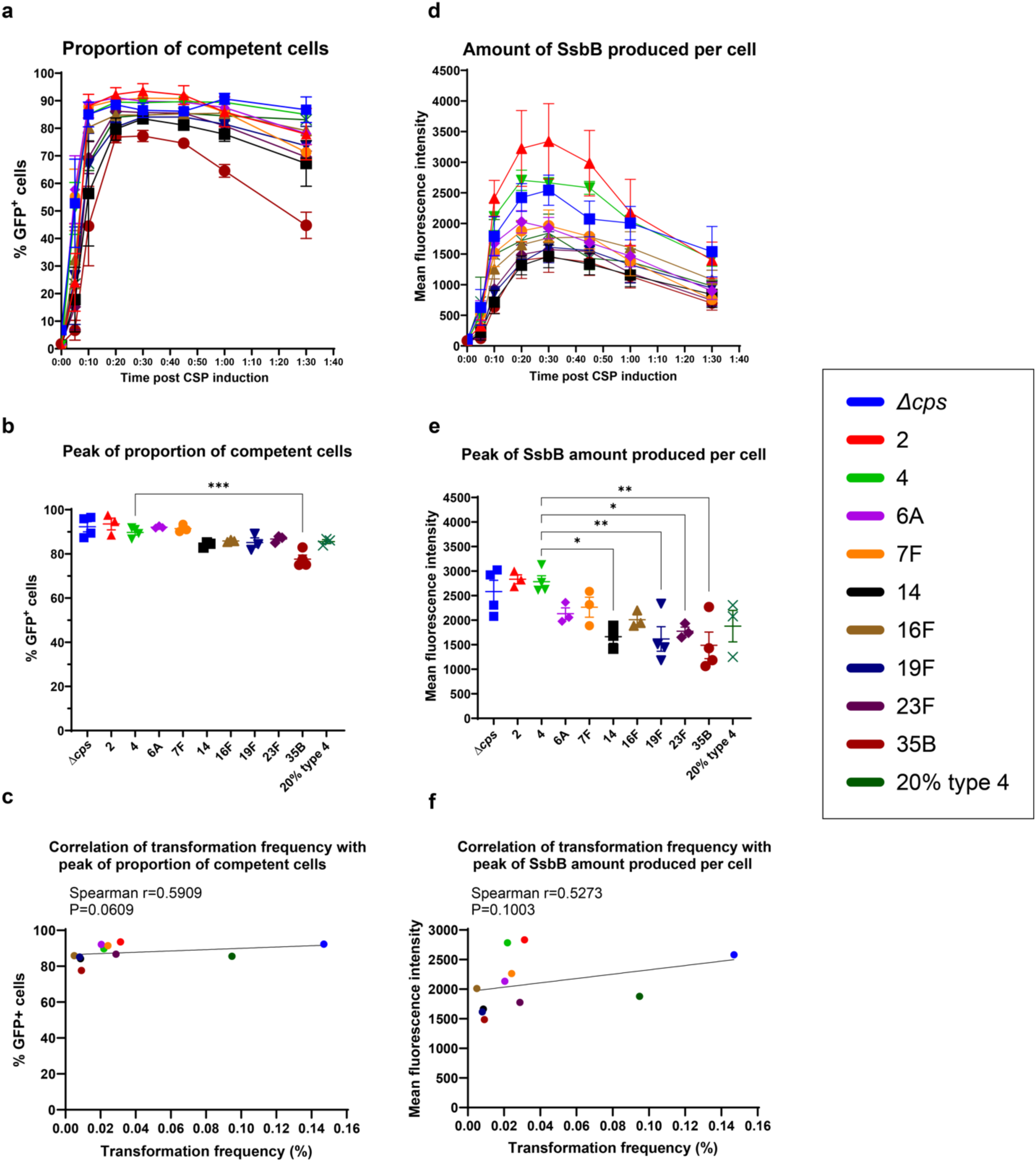
Competence level of isogenic capsule-switch strains examined by P*_ssbB_- ssbB-gfp* reporters. (a) Kinetics of proportion of competent cells post CSP induction. The percentage of GFP+ cells, which represent competent cells, was examined by flow cytometry at selected time points post CSP induction. (b) Peak value of proportion of competent cells after CSP induction. Statistical significance was determined using an ordinary one-way ANOVA with Tukey’s multiple comparisons test. For brevity, only comparison to type 4 with significance was shown on the graph. (c) Correlation of transformation frequency with PCR product and the peak value of proportion of competent cells. Spearman nonparametric correlation coefficients were computed. (d) Kinetics of amount of SsbB produced per cell post CSP induction. Mean fluorescence intensity, which represents amount of SsbB produced per cell, was examined by flow cytometry at selected time points post CSP induction. (e) Peak value of amount of SsbB produced per cell after CSP induction. Statistical significance was determined using an ordinary one-way ANOVA with Tukey’s multiple comparisons test. For brevity, only comparisons to type 4 with significance were shown on the graph. (f) Correlation of transformation frequency with PCR product and the peak value of amount of SsbB produced per cell. Spearman nonparametric correlation coefficients were computed. Strains are designated by their capsule type or amount for brevity. Data show mean ± SEM from at least 3 independent experiments. *, P≤0.05; **, P≤0.01; ***, P≤0.001.

### Capsule impedes Com pilus surface accessibility

We then evaluated the influence of capsule on the DNA uptake process. As the central component of the transformation apparatus, Com pilus can extend up to 2-3 µm beyond the cell surface to bind and take in extracellular DNA [16]. We postulated that the shielding effect of capsule might inhibit the exposure of Com pilus at the cell surface. The backbone of Com pilus consists of the major pilin, ComGC, assembled into the growing filament during pilus extension from a preformed pool of subunit monomers in the cell membrane [16, 20, 35]. We first compared ComGC abundance during competence in the *cps*- strain and the strain producing native type 4 capsule via Western blot analysis. ComGC production, which peaked at 15 min post-induction, was exclusively detected in competent cells in both strains (Fig. 3a). Densitometry analysis of immunoblots revealed similar levels of ComGC in the cell fraction (Fig. 3b). The amount of ComGC in supernatant, likely a reflection of pilus shearing during sample preparation, was significantly higher in the *cps*- compared to the type 4 strain (Fig. 3b), suggesting the *cps*- strain collectively produced more ComGC during competence.

**Fig 3.**
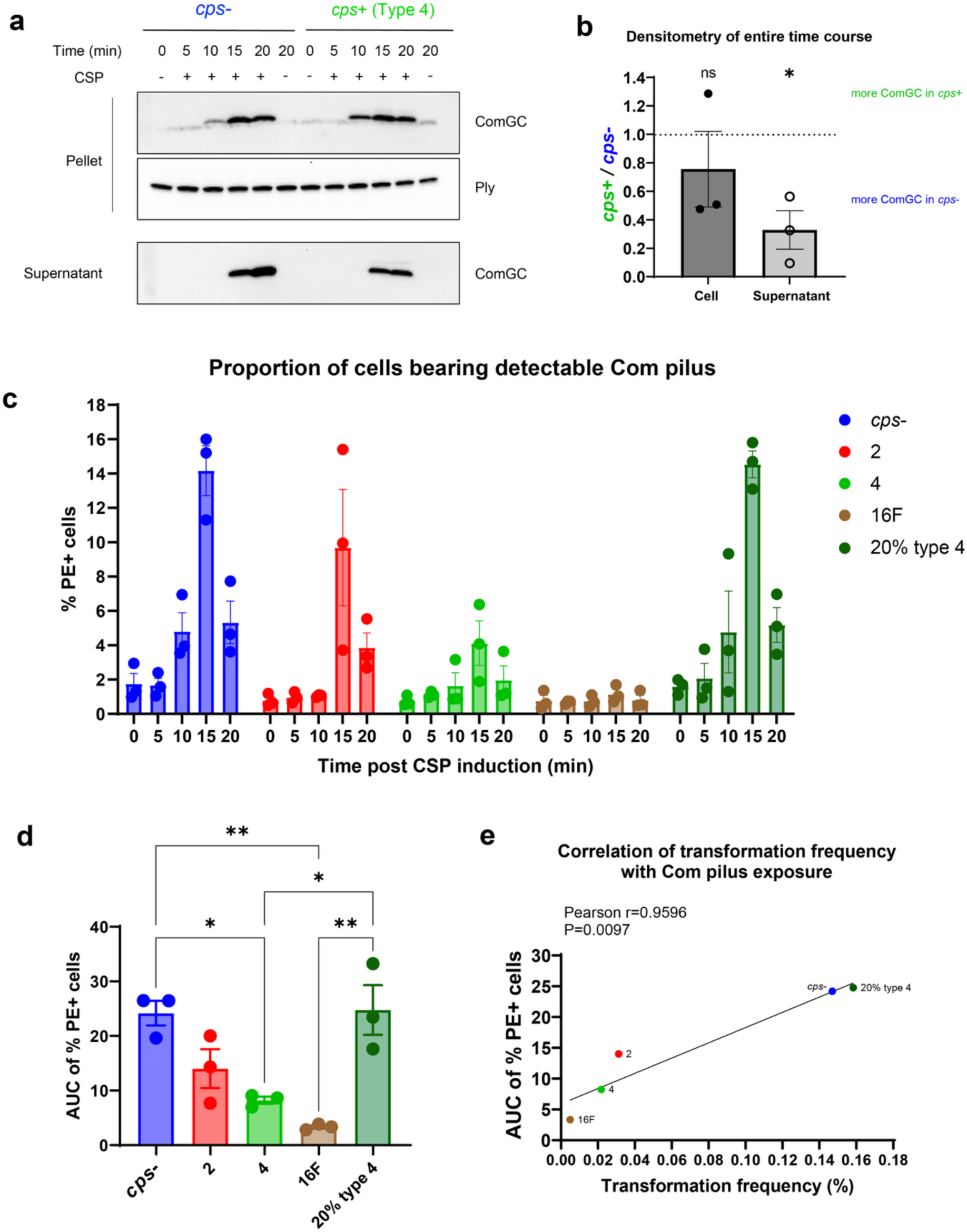
Production of ComGC and exposure of Com pili at the cell surface. (a) Representative immunoblots of ComGC associated with the bacterial cells or in the culture supernatant in *cps*- or type 4 strains at different time points post CSP induction. Pneumolysin (Ply) was used as a loading control for bacterial cell fraction. Sample loading was standardized by equalizing total protein amounts obtained from the cell pellet sample collected at 0 min post-induction. Loading of supernatant samples was proportional to their corresponding bacterial cell pellet samples to ensure equivalent representation of the original culture volume prior to centrifugation. (b) Quantification of ComGC signal associated with the bacterial cell or in the culture supernatant from the immunoblot results. At 10, 15, and 20 min post CSP induction, respectively, the density of ComGC signal of type 4 strain was compared to that of *cps*-. The average of these ratios was plotted on the graph to reflect the average level of the entire time course, and compared to a hypothetical mean of 1 using the one sample t test. (c) Kinetics of proportion of cells bearing Com pilus detectable by ComGC antibody post CSP induction. At selected time points post CSP induction, bacterial cells were stained with ComGC antibody and PE-conjugated secondary antibody, and the percentage of PE+ cells was examined by flow cytometry. (d) Area under the curve (AUC) of the kinetics shown in (c). AUC of kinetics from each independent experiment was calculated. Statistical significance was determined using an ordinary one-way ANOVA with Tukey’s multiple comparisons test. (e) Correlation of transformation frequency with PCR product and AUC of proportion of cells bearing detectable Com pilus. Pearson correlation coefficients were computed. Strains are designated by their capsule type or amount for brevity. Data show mean ± SEM from 3 independent experiments. ns, not significant; *, P≤0.05; **, P≤0.01.

Therefore, we then investigated if capsule affects the exposure of Com pilus at the cell surface by examining the kinetics of its accessibility in fixed, competent cells by flow cytometry with an antibody to ComGC. We assessed five representative constructs with different types or amounts of capsule that exhibited high, medium, or low transformation frequencies. Pilus was detected earlier (10 min post-induction) in the *cps*- and 20% type 4 capsule strains, which had the highest transformation frequency (Fig. 3c). This disparity suggests that full encapsulation hinders pilus extrusion during initial assembly. Quantification by area under the curve (AUC) of the exposure kinetics over 20 minutes revealed significantly higher cumulative pilus exposure in the populations of *cps*- and 20% type 4 strains compared to native type 4 and type 16F strains (Fig. 3d). Strikingly, AUC values strongly correlated with transformation frequency (Fig. 3e), suggesting a relationship between surface exposure of Com pilus and transformation efficiency. Further analyses demonstrated that neither the proportion of competent cells nor the SsbB amount per cell of the five representative strains correlated with their transformation frequency (Fig. S2a and S2b), reinforcing that variations in transformability were specifically due to differences in Com pilus surface exposure rather than the level of competence.

Regarding the effect of capsule amount, the strain producing 20% type 4 capsule closely resembled the *cps*- strain in both transformation efficiency and Com pilus exposure, contrasting sharply with the strain producing native type 4 capsule (Fig. 1b, 1c, 3c, and 3d). Transmission electron microscopy (TEM) revealed sparse, incomplete capsule coverage of the cell surface in 20% type 4 strain (Fig. S3), suggesting that capsule must fully surround the cell to impede pilus assembly. Collectively, these results demonstrate that capsule imposes a serotype- and quantity-dependent physical barrier to hinder pilus extrusion.

### Capsule restricts Com pilus prevalence and stability

To further characterize Com pilus surface exposure in capsule-switch strains, we directly visualized Com pili using TEM. Since the parent TIGR4 strain also produces other types of pili, we used immunogold labeling to specifically visualize Com pili at 15 min post CSP induction [36, 37]. Considering the strain producing 20% type 4 capsule resembled the *cps*- strain, we just included the *cps*- strain along with type 2, type 4, and type 16F strains. Since the capsule is a highly hydrated structure which collapses during water removal in the TEM sample preparation process, we were unable to visualize capsule under this experimental condition. In each construct, we could observe specifically labeled Com pili attached to the cell surface (Fig. 4a), as well as Com pili sheared from the cell in the background. Quantitative analysis indicated the *cps*- strain exhibited a significantly higher frequency of cells bearing Com pilus compared to all encapsulated strains (Table 3). There were serotype-specific effects among encapsulated strains as well, as the proportion of piliated cells in type 2 and type 16F strains were both significantly different from that in the native type 4 strain (Table 3). Importantly, a strong positive correlation was observed between transformation frequencies and the proportion of cells bearing Com pilus (Fig. 4b), suggesting that the frequency of Com pilus-bearing cells directly determines transformation efficiency.

**Table 2.**
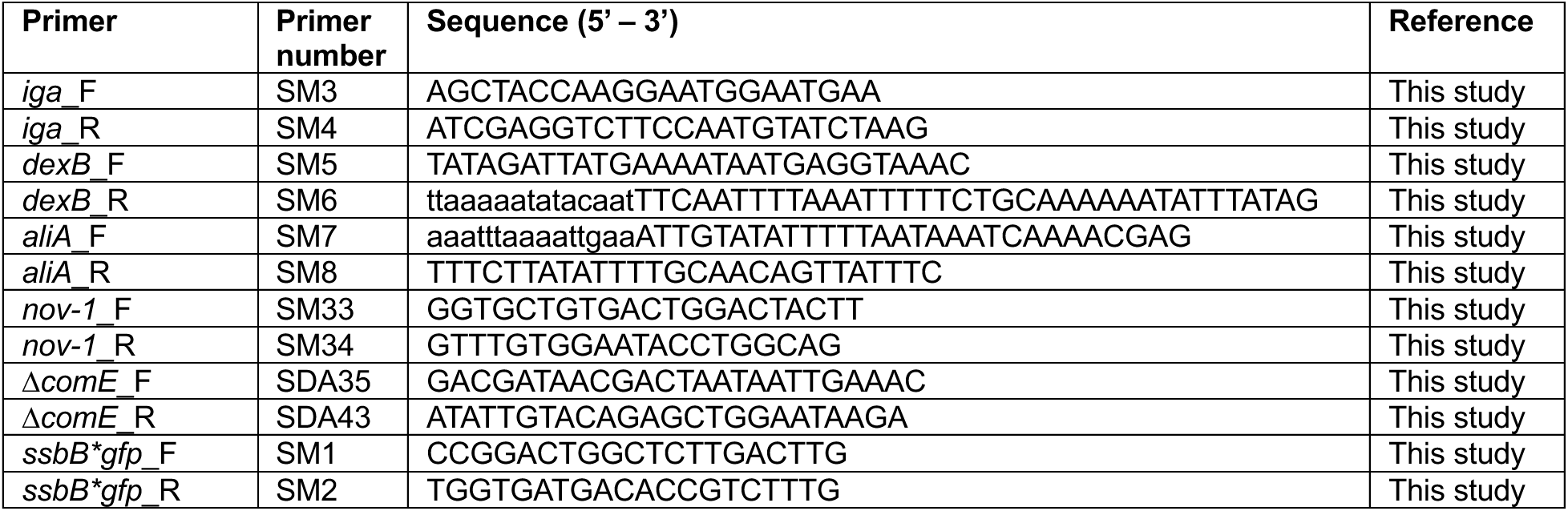
Primers used in this experimental work.

**Table 3.** Frequency of Com pili visualized by transmission electron microscopy.

| Strain capsule type | Number of cells screened | Total Com pili |  | Cells bearing Com pilus |  | Frequency of sheared Com pili<sup>c</sup> | Proportion of cells bearing Com pilus |  | Proportion of sheared Com pili |
| --- | --- | --- | --- | --- | --- | --- | --- | --- | --- |
|  |  | Number | Frequency<sup>a</sup> | Number | Frequency<sup>b</sup> |  | P value compared to *cps*-<sup>d</sup> | P value compared to 4<sup>e</sup> | P value compared to *cps*-<sup>f</sup> |
| *cps*- | 637 | 210 | 32.97% | 178 | 27.94% | 15.24% | - | <0.0001 | - |
| 2 | 823 | 69 | 8.38% | 43 | 5.22% | 37.68% | <0.0001 | 0.0303 | 0.0001 |
| 4 | 967 | 69 | 7.14% | 30 | 3.10% | 56.52% | <0.0001 | - | <0.0001 |
| 16F | 778 | 8 | 1.03% | 1 | 0.13% | 87.50% | <0.0001 | <0.0001 | <0.0001 |

Competent bacterial cells were stained with ComGC antibody and protein A conjugated to 10-nm gold particles and examined by transmission electron microscopy. There were both cell-associated Com pili and sheared Com pili at the background.
a: Calculated as the number of total Com pili in the view divided by the number of cells screened.
b: Calculated as the number of cells bearing Com pilus divided by the number of cells screened.
c: Calculated as the number of sheared Com pili divided by the number of total Com pili.
d: Calculated using Fisher's exact test to compare the proportion of cells bearing Com pilus of each strain to that of *cps*- strain.
e: Calculated using Fisher's exact test to compare the proportion of cells bearing Com pilus of each strain to that of type 4 strain.
f: Calculated using Fisher's exact test to compare the proportion of sheared Com pili of each strain to that of *cps*- strain.

**Fig 4.**
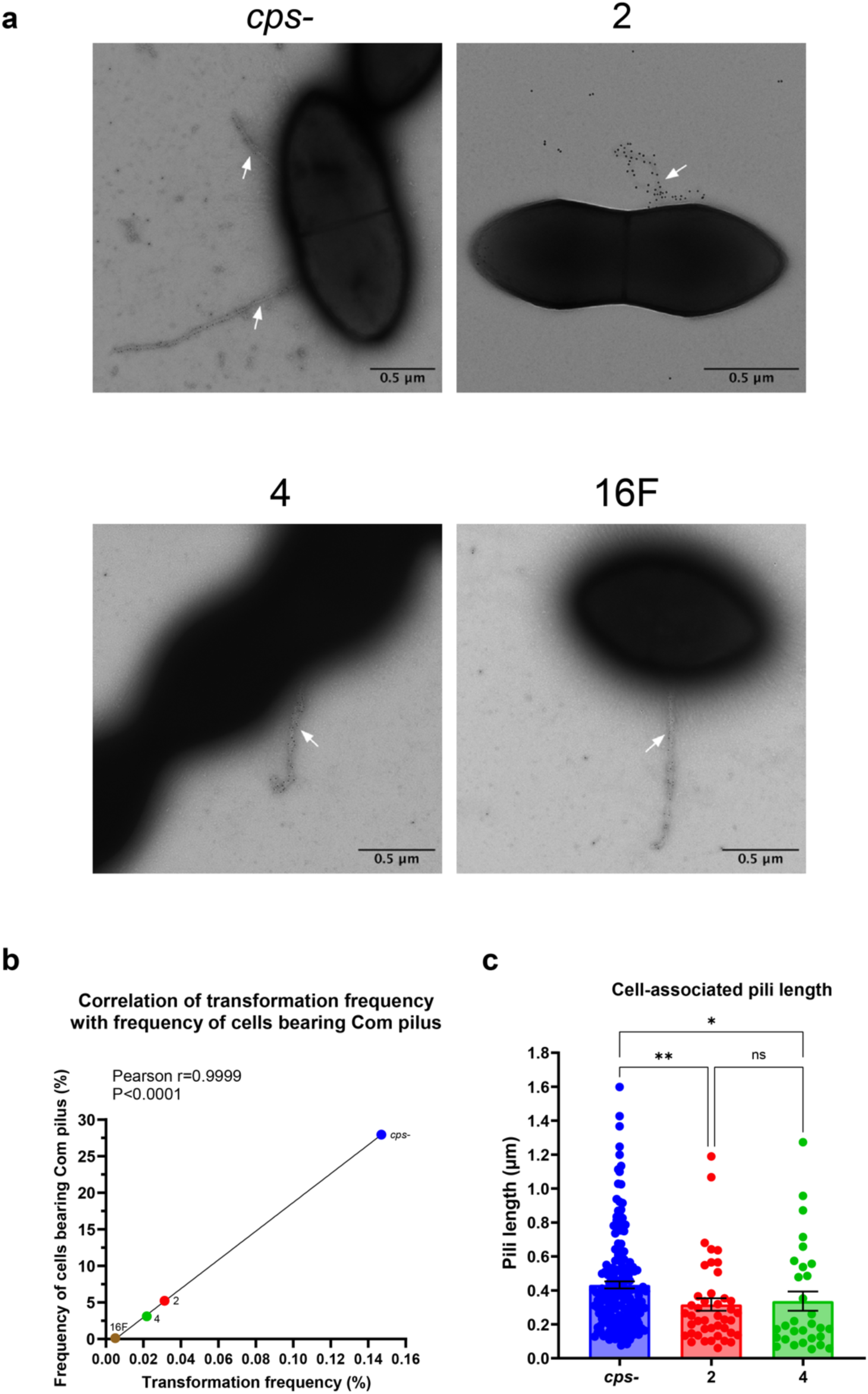
Frequency and length of cell-associated Com pili examined by transmission electron microscopy. (a) Representative immunogold electron micrographs to visualize Com pili in different strains. Competent bacterial cells were stained with ComGC antibody and protein A coupled to 10-nm gold particles. White arrows indicate the Com pilus. Scale bar: 0.5 µm. (b) Correlation of transformation frequency with PCR product and frequency of cells bearing Com pilus visualized by electron microscopy. Pearson correlation coefficients were computed. (c) Length of cell-associated Com pili. Data show mean ± SEM from measurement of at least 30 Com pili. Type 16F strain was excluded due to inadequate pilus counts. Statistical significance was determined using Kruskal-Wallis test with Dunn’s multiple comparisons test. ns, not significant; *, P≤0.05; **, P≤0.01. Strains are designated by their capsule type for brevity.

Furthermore, pilus length measurements indicated that the *cps*- strain produced significantly longer Com pili compared to type 2 and type 4 strains (Fig. 4c). This finding suggests that capsule presence also impedes Com pilus elongation, potentially limiting DNA-binding capacity and thereby reducing transformation efficiency in encapsulated strains.

Interestingly, the *cps*- strain exhibited a significantly lower frequency of sheared Com pili compared to all encapsulated strains, while type 16F strain, which had the lowest transformation frequency, exhibited the highest proportion of sheared Com pili (Fig. 1b and Table 3). This result implies increased pilus fragility in encapsulated strains, exacerbating transformation deficits by reducing functional pilus stability.

### Relationship between capsule characteristics and transformation frequency

We explored how distinct characteristics of each serotype contributed to different inhibitory effects on transformation. We first questioned whether differences in capsule thickness across serotypes impose distinct hindrance on pilus biogenesis, potentially driving disparities in the presence and length of Com pili. We quantified capsule thickness of each capsule-switch strain using fluorescence light microscopy with a modified dextran exclusion assay, where capsule thickness is determined as the shadow extending from the Nile red-labeled cell surface to the edge of FITC-dextran exclusion (Fig. S4) [38]. Fully encapsulated strains exhibited significant variation in capsule thickness, with serotypes 16F, 19F, and 23F producing thicker capsules (Fig. 5a). However, capsule thickness did not correlate with transformation frequency (Fig. 5b), indicating that factors other than thickness, such as specific polysaccharide composition or structural characteristics of the capsule, could significantly influence the inhibitory effect on transformation. In addition, we also examined the number of carbons and high energy bonds per polysaccharide repeat unit, which have been suggested to correlate with capsule thickness, and found neither correlated with transformability (Fig. S5a and S5b) [39].

**Fig 5.**
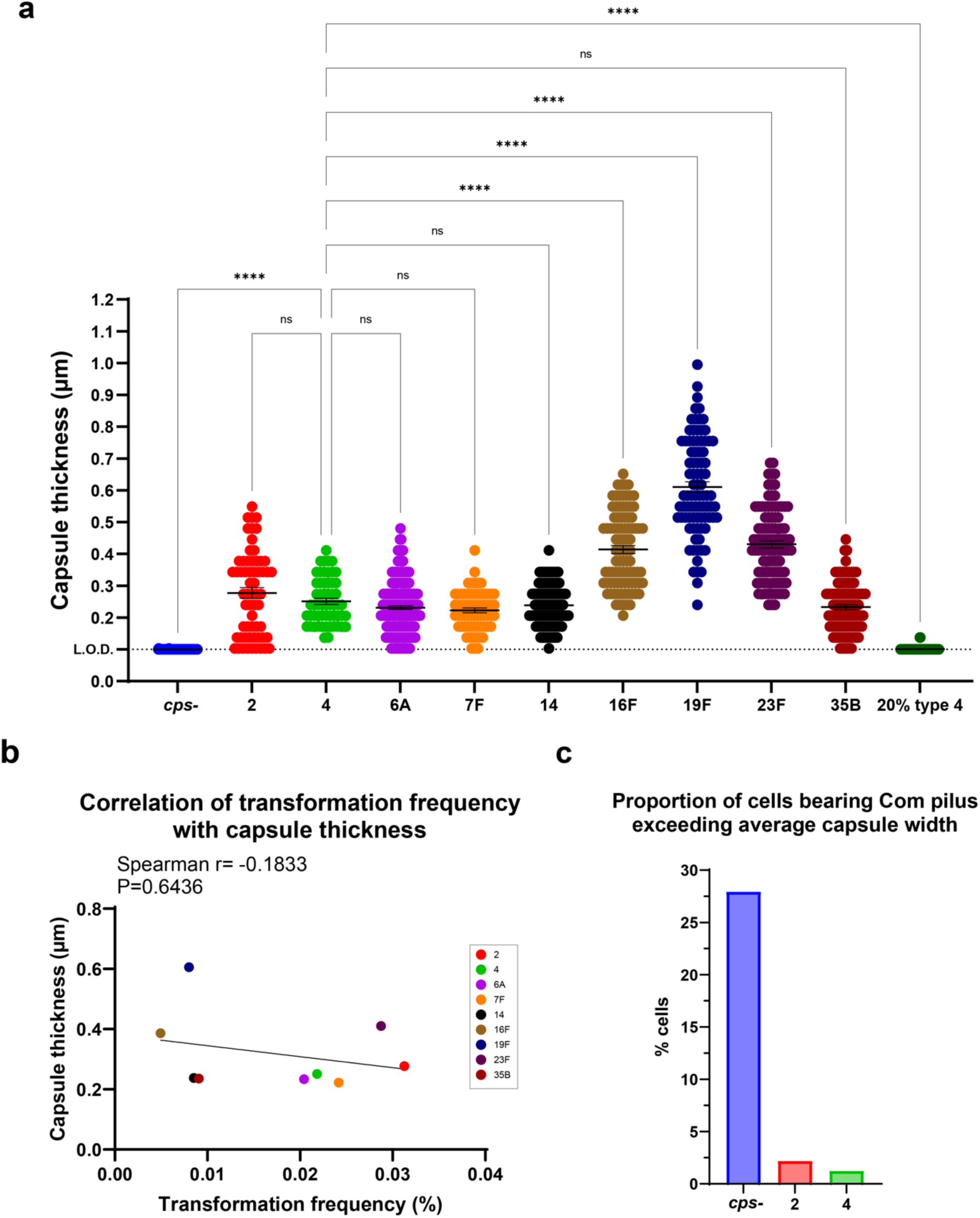
Capsule thickness and comparison to the length of Com pili. (a) Capsule thickness as determined by the distance between bacterial surface and the edge of the shadow excluded by FITC-dextran (M.W. 2,000 kDa) visualized by light microscopy. Dashed line represents limit of detection. Data show mean ± SEM from measurement of at least 56 individual cells of varying morphology. Statistical significance was determined using Kruskal-Wallis test with Dunn’s multiple comparisons test. For brevity, only comparisons to type 4 were shown on the graph. ns, not significant; ****, P≤0.0001. (b) Correlation of transformation frequency with PCR product and capsule thickness. Spearman nonparametric correlation coefficients were computed. (c) Proportion of cells with Com pilus that exceeds average capsule thickness. The average capsule thickness of type 2 and type 4 were calculated from (a); the capsule thickness of *cps*- was considered as 0. The number of cells with a Com pilus that exceeds average capsule thickness was determined from Fig. 4c. It was then divided by the number of total cells screened as in Table 3. Strains are designated by their capsule type or amount for brevity.

Based on the quantification of capsule thickness and Com pili length (Fig. 4c and 5a), we assorted the proportion of cells that formed a Com pilus exceeding the average capsule width in each serotype. Compared to the *cps*- strain, markedly fewer cells in the type 2 and type 4 encapsulated strains displayed pili long enough to extend beyond the capsule boundary (Fig. 5c). This parameter could potentially represent a more accurate predictor of transformability, assuming effective DNA uptake requires Com pili length to surpass capsule thickness.

Next, we assessed the impact of capsule porosity on transformation by evaluating the accessibility of FITC-dextran molecules of various sizes to the bacterial surface. Each capsule-switch strain was stained with 4, 20, 40, 60 (60-76), and 150 kDa FITC-dextran, respectively, and the exclusion zone from the cell membrane to FITC-stained background was visualized and quantified via fluorescence microscopy [38]. The threshold of exclusion was determined as the smallest dextran size at which the exclusion width significantly differed from the 4 kDa negative control. Serotypes 2, 4, 6A, 7F, and the 20% type 4 strain, which were highly transformable strains, had more porous capsules with larger thresholds of exclusion (60-150 kDa) (Fig. 6). In contrast, serotypes 16F, 19F, and 35B, having lower transformation frequencies, showed less porous capsules with smaller thresholds of exclusion (20-40 kDa) (Fig. 6). Serotypes 14 and 23F displayed intermediate thresholds (40-60 kDa) (Fig. 6). These findings suggest that capsule porosity contributes substantially to transformation inhibition, with less porous capsules imposing greater steric hindrance on Com pilus assembly and function.

**Fig 6.**
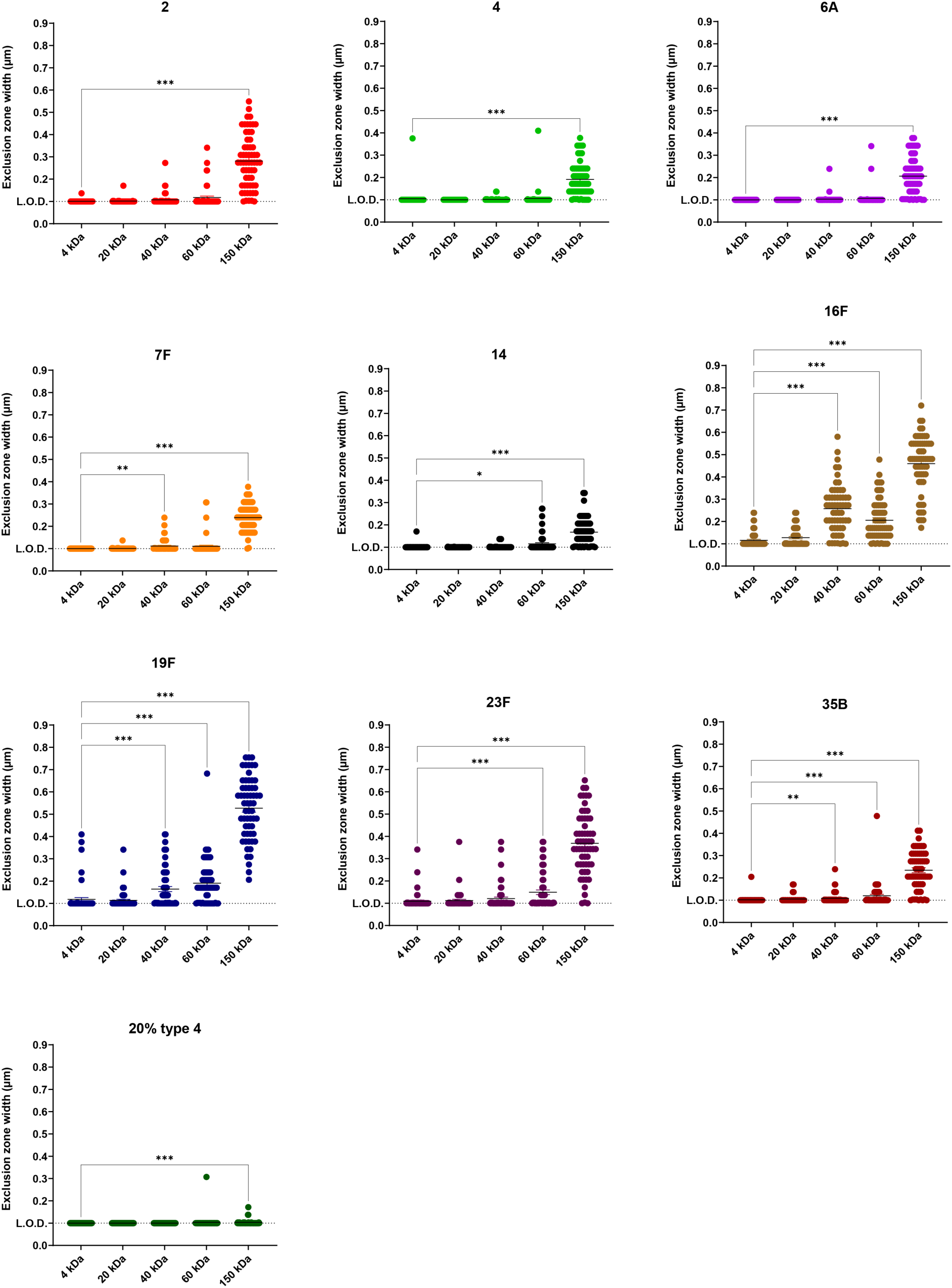
Capsule porosity of each serotype. Access of varying sizes of FITC-dextran to the bacterial surface was visualized by light microscopy. The width of exclusion zone by capsule was determined as the distance between bacterial cell membrane and the edge of the shadow by excluded FITC-dextran. Dashed line represents limit of detection. Data show mean ± SEM from measurement of 60 individual cells of varying morphology per condition. Each dextran size group was compared to the 4 kDa group using Kruskal-Wallis test with Dunn’s multiple comparisons test. *, P≤0.05; **, P≤0.01; ***, P≤0.001. Strains are designated above by their capsule type or amount for brevity.

## DISCUSSION

Despite the paradigm of transformation developed using unencapsulated laboratory *Spn* strains, understanding of transformability of clinically relevant encapsulated strains is limited. It is well established that within laboratory environment, spontaneous variations in colony morphology occur naturally across serotypes, with opaque colonies producing more capsule and exhibiting lower transformation rates compared to less encapsulated transparent colonies [25, 40]. However, opacity differences are governed by genome-wide differential DNA methylation due to DNA inversions in homologous *hsdS* genes, confounding a direct systematic assessment of how capsule amount alone affects transformation [41]. Here, we utilized isogenic strains differing exclusively at the *cps* locus to specifically isolate the impact of capsule on transformation. We demonstrated that the presence of a capsule sufficiently thick to fully envelop the cell significantly inhibits transformation through impeding Com pilus assembly on the cell surface. Further analysis of isogenic strains with diverse serotypes revealed serotype-specific suppression on Com pilus assembly and retention and thus transformability, with less porous capsules showing the greatest inhibition.

Among nine tested serotypes, we specifically examined serotypes 16F and 35B, since they were previously reported to exhibit extremely low or high recombination rates during human carriage [26]. Surprisingly, serotype 35B showed a low transformation frequency *in vitro* (Fig. 1b and 1c). This discrepancy likely arises from prolonged carriage duration in natural settings, where thicker capsules confer survival advantages and increase opportunities for recombination—factors not replicated under controlled uniform experimental conditions [26]. Additionally, clinical isolates possess heterogeneous genetic backgrounds, introducing confounding variables beyond serotype alone. Therefore, although prior work suggested that recombination rate increases with capsule size, such rates represent cumulative phenotypes shaped by multifactorial interactions. It is also worth noting that the capsule thickness data in this prior work were derived from another study using the same serotypes but not same lineages [39], potentially causing discrepancies of actual capsule sizes due to substantial intra-serotype variation in capsule production, as we recently reported [38].

Western blot analysis of ComGC in culture supernatant indicated extensive pili shearing by sample centrifugation. Enhanced shearing in the *cps*- strain likely reflects increased surface exposure and assembly frequency of Com pili compared to the type 4 strain (Fig. 3a-d, Table 3). In contrast, the comparable amount of cell-associated ComGC in both strains were likely to be predominantly unassembled subunits (Fig. 3b). Therefore, cultures were prefixed with paraformaldehyde prior to centrifugation to minimize pili shearing for subsequent Com pilus exposure assay and TEM. Interestingly, TEM of prefixed samples revealed higher pilus fragility in encapsulated strains compared to *cps*- (Table 3), possibly due to interactions between capsular polysaccharides and pilin subunits that alters pilus assembly dynamics.

These observations highlight the necessity for future research addressing how distinct capsular architectures modulate Com pilus assembly and functionality. Although capsule-mediated inhibition of transformation is serotype- and quantity-dependent, and capsule-switch strains exhibit varying capsule thicknesses, transformation efficiency disparities among these strains cannot be attributed solely to capsule thickness (Fig. 5b). Prior studies have proposed associations between capsule thickness and compositional metrics such as carbon count or ATP-equivalent energy expenditure per polysaccharide repeat unit [39]. Neither of these metrics, however, correlated with transformation frequency (Fig. S5a and S5b). Additionally, capsule surface charge does not significantly affect transformation efficiency, as serotypes 7F and 14—the only two serotypes with neutral charge—did not show a distinct transformation pattern compared to the other serotypes with negatively charged capsules (Fig. 1b and 1c). These results implicate additional capsule-specific physicochemical properties in mediating transformation inhibition. Indeed, our results identified capsule porosity as a critical determinant shaping the capsule’s inhibitory effect on transformation (Fig. 6). We propose that structural variations among polysaccharides modulate capsule porosity, where more porous capsules permit steric accommodation of ComGC subunit extrusion, thus correlating with higher transformability. Additional serotype-specific chemical and structural properties, including spatial conformation, branching patterns, or intermolecular interactions, may also collectively shape the interaction of polysaccharide with Com pilus through mechanisms yet to be elucidated.

This study advances our understanding of pathogenic *Spn* strains and has critical implications for pneumococcal biology and pathogenesis research. Since there are natural variations of capsule production within cell populations (Fig. 5a), and capsule amount affects transformation, there may be selection biases during mutant construction through transformation, favoring hyper-transformable colonies which are less encapsulated. Such biases must be carefully considered when assessing virulence of engineered strains, since capsule quantity critically influences pathogenic potential. Moreover, our results reveal how the relationship between capsule and transformation affects the persistence and prevalence of different serotypes and underscores evolutionary trade-offs of the bacterium as a human pathogen. *Spn* evolves highly heterogenous serotypes under immune pressures, but *Spn* carriage is characterized by the predominance of a limited array of serotypes. While some types of capsule may shield the cell surface and enhance survival in hostile host environments in the short term, they may not favor transformation and thus limit access to genetic materials with new traits, constraining long-term adaptability and fitness. The prevalence of certain serotypes likely reflects a balance between capsule’s protective function and the genetic flexibility afforded by high transformability. Capsules that not only effectively shield the cell surface against host attack but also allow transformation should be fit *in vivo* and also genetically adaptable to environmental pressures (e.g. antibiotics, immune pressures) for the long term. In addition, the existence of naturally non-encapsulated strains suggests another intriguing strategy *Spn* may employ to reconcile virulence and adaptation [42]. These non-encapsulated strains commonly colonize humans asymptomatically and may serve as a gene reservoir to disseminate virulence genes through elevated transformability. These non-encapsulated strains could also revert to encapsulated to evade host defense and retain stable colonization. Collectively, these findings suggest the recombination and adaptation of *Spn* within the host involve a complex interplay of factors that cooperatively contribute to the ultimate evolutionary fitness of the species.

## MATERIALS AND METHODS

### Bacterial growth and culture conditions

*Spn* strains used in this study are listed in Table 1. Unless otherwise specified, strains were grown statically at 37°C in Todd Hewitt broth (Becton Dickinson) supplemented with 1% yeast extract (Thermo Fisher Scientific) (THY) at pH 6.6 to suppress endogenous competence development. Bacterial colonies from tryptic soy (TS) agar plates containing 5% sheep blood (Becton Dickinson) were inoculated into 5 mL THY and grown to early exponential phase. The culture was sub-cultured into fresh medium to an initial optical density (OD) of 0.005 at 600 nm and harvested at OD of 0.05 for competence induction and subsequent analyses.

For quantification of colony-forming unit (CFU) and transformant selection, unless otherwise specified, the culture was serially diluted in phosphate-buffered saline (PBS) and plated on TS agar supplemented with catalase (3,700 U/plate; Worthington Biochemical Corporation). Plates were incubated overnight at 37°C in 5% CO_2_.

### Bacterial strain construction

Primers used in strain construction are listed in Table 2. The construction of isogenic capsule-switch strains in isolate TIGR4 (T4) genetic background was previously described [27]. Briefly, an unencapsulated mutant T4*^cps::Sweet^ ^Janus^* (P2422) was transformed with genomic DNA from *Spn* clinical isolates expressing type 2, 4, 6A, 7F, 14, 16F, 19F, 23F, or 35B capsules, followed by selection on TS agar plates supplemented with streptomycin (200 µg/mL) and 10% sucrose [43]. Smooth colonies were verified for desired serotypes via colony immunoblot and backcrossed twice with P2422 to minimize background mutations. The T4^20%^ ^CPS^ strain was constructed by replacing the native *cps* promoter with the *cps* promoter from serotype 23F [27].

P2742 was generated by transforming P2422 with plasmid pE539 containing a spectinomycin resistance (Spec^R^) cassette [44]. Transformants were selected on TS agar plates containing spectinomycin (200 µg/mL) and confirmed via PCR using primers SM3/SM4.

P2746 was constructed to delete the Sweet Janus cassette from the *cps* locus in P2422. Genes flanking the *cps* locus, *dexB* (upstream) and *aliA* (downstream), were amplified using primers SM5/SM6 and SM7/SM8, respectively. Overlapping regions in primers SM6 and SM7 facilitated ligation of the *dexB* and *aliA* PCR products using the NEBuilder HiFi DNA Assembly kit (New England Biolabs). P2422 was transformed with the assembled *dexB*-*aliA* cassette and selected on TS agar plates containing streptomycin (200 µg/mL) and screened for kanamycin (250 µg/mL) sensitivity. Selected transformants were confirmed by PCR with primers SM5/SM8.

Strains used in cell-to-cell gene transfer assay were generated by transforming P2746 (unencapsulated) or P1542 (encapsulated) with either the *nov-1* or *comE::Janus* PCR products. The *nov-1* fragment, amplified from strain P1281 with primers SM33/SM34, conferred novobiocin resistance; whereas *comE::Janus* amplified from strain P2074 with primers SDA35/SDA43 conferred kanamycin resistance and rendered the recipient competence-deficient. Transformants were selected on TS agar plates containing novobiocin (10 µg/mL) or kanamycin (250 µg/mL), respectively, and *comE::Janus* mutants were confirmed by PCR and by demonstrating transformation deficiency.

Each capsule-switch strain was transformed with the P*_ssbB_-ssbB-gfp* cassette to generate a GFP reporter for competence development. The cassette was amplified from strain P2670 using primers SM1/SM2. Transformants were selected on TS agar plates containing kanamycin (250 µg/mL) and verified by PCR.

### Transformation

Recipient strains were cultured as described above to an OD_600_ of 0.05 in THY (pH=6.6). Unless otherwise specified, cultures were incubated with 150 ng/mL CSP-II (Eurogentec) and 300 ng/mL Spec^R^ PCR product for 30 min at 37°C (conditions were determined based on pilot experiments for high yield). Transformed cultures were serially diluted in PBS and plated on TS agar plates containing spectinomycin (200 µg/mL) for transformants selection, or on plain TS agar plates for total counts. Transformation frequency was calculated as the density of transformants (CFU/mL) divided by the total cell density (CFU/mL).

For the transformation with mixed culture of seven serotypes, each strain was individually cultured to an OD_600_ of 0.05 and mixed in equal proportions. The combined culture was transformed with the Spec^R^ PCR product and plated as described above. Colonies from both the transformant and total population were transferred to nitrocellulose membranes for serotype identification by immunoblot as previously described [45]. The number of *cps*- colonies was determined by subtracting the count of colonies positive for the other six serotypes from the total colony number.

### Cell-to-cell gene transfer assay

Strains cultured to an OD_600_ of 0.05 were concentrated 20-fold in fresh THY (neutral pH). Concentrated attacker and victim cultures were mixed 1:1 and incubated with an equal volume of inducer cocktail for 1 h at 37°C. The inducer cocktail was freshly prepared before experiment as described previously, with modifications to include 1 µg/mL CSP-II, 0.008% BSA, and 1 mM CaCl_2_ in THY [18]. The culture was serially diluted in fresh THY and plated in soft agar as previously described, with the fourth layer containing novobiocin (10 μg/mL) and kanamycin (250 μg/mL) for selection of dual-resistant colonies, or plain THY agar for total colony counts [46]. The efficiency of cell-to-cell gene transfer was calculated as the density of dual-resistant colonies (CFU/mL) divided by the total colony density (CFU/mL).

### Competence development assay

Strains harboring the P*_ssbB_-ssbB-gfp* reporter were cultured to an OD_600_ of 0.05 and induced with 150 ng/mL CSP-II at 37°C. At 0, 5, 10, 20, 30, 45, 60, and 90 min post-induction, 1 mL aliquots were collected, immediately centrifuged at 10,000 g for 1 min at 4°C, and resuspended in PBS containing 1% paraformaldehyde (PFA) to be fixed for 30 min at 4°C. After washing with DPBS, cells were resuspended in 150 µL flow cytometry buffer (DPBS containing 0.5 mM EDTA) for analyses. Uninduced controls were processed identically. Flow cytometry was performed on an LSRII flow cytometer (Becton Dickinson) using BD FACSDiva software (Becton Dickinson) for data acquisition, and data were analyzed using FlowJo software (Tree Star).

### Western blot

Strains cultured to an OD_600_ of 0.05 were induced with 150 ng/mL CSP-II at 37°C. At 0, 5, 10, 15, and 20 min post-induction, 2 mL aliquots were collected and immediately centrifuged at 2,000 g for 4 min at 4°C. Cell pellets were mixed with lysing buffer (1% Triton X-100 in PBS), Tris-glycine SDS sample buffer (Thermo Fisher Scientific), and NuPAGE sample reducing agent (Thermo Fisher Scientific), followed by heating at 85°C for 2 min. Supernatants were filtered through a 0.2 µm filter and precipitated at 4°C overnight with 10% trichloroacetic acid. Precipitated proteins were pelleted, washed with acetone, solubilized in 8 M urea, and mixed with SDS sample buffer and reducing agent with heating. Uninduced controls were processed identically.

The immunoblot was performed as described previously with a modified protocol [47]. Samples were separated using Novex 10-20% Tris-Glycine Plus WedgeWell Gels (Thermo Fisher Scientific) and transferred onto PVDF membranes using the iBlot3 (Thermo Fisher Scientific). Membranes were probed with rabbit α-ComGC_Δ1-39_ (provided by Birgitta Henriques-Normark) at 1:3,000 dilution, followed by incubation with HRP-conjugated goat anti-rabbit IgG (Thermo Fisher Scientific) at 1:10,000 dilution and development with SuperSignal West Pico PLUS Chemiluminescent Substrate (Thermo Fisher Scientific) [17]. For pneumolysin detection, membranes were probed as described [47]. Immunoblots were imaged, and densitometry was performed using the iBright Imaging System (Thermo Fisher Scientific).

### Com pilus exposure assay

Strains cultured to an OD_600_ of 0.05 were induced with 150 ng/mL CSP-II at 37°C. At 0, 5, 10, 15, and 20 min post-induction, 3 mL aliquots were collected and fixed with 1% PFA for 30 min. Cells were pelleted at 2,000 g for 4 min and gently resuspended in 200 µL flow cytometry buffer. Half of each sample (100 µL) was stained with rabbit α-ComGC_Δ1-39_ at a dilution of 1:200 for 30 min, followed by a DPBS wash. Subsequently, cells were stained with R-phycoerythrin (PE)-conjugated goat anti-rabbit IgG (Jackson ImmunoResearch) diluted 1:1,000 for 30 min and washed. The remaining 100 µL served as a gating control, stained only with the secondary antibody. Stained cells were gently resuspended in 100 µL ice-cold flow cytometry buffer for analyses. All steps were carried out at 4°C. Flow cytometry was performed on a BD FACSymphony A5 cell analyzer (Becton Dickinson) using BD FACSDiva software (Becton Dickinson) for data acquisition, and data were analyzed using FlowJo software (Tree Star).

### Transmission electron microscopy

For imaging of immunogold-labeled Com pilus, strains were cultured to an OD_600_ of 0.05 and induced with 150 ng/mL CSP-II for 15 min at 37°C. Following fixation with 1% PFA for 30 min at 4°C, cells were pelleted at 2,000 g for 4 min at 4°C and gently resuspended in 50 µL fresh THY. Aliquots of 10 µL were placed on parafilm for immunogold labeling. Formvar carbon-coated copper grids (Electron Microscopy Sciences) were glow-discharged for 30 sec and placed face down on the bacterial samples for 2-5 min. Grids were fixed with 10 µL of 0.2% glutaraldehyde in PBS for 2 min, washed thrice with PBS, and incubated with 10 µL of 50 mM glycine for 5 min to quench residual fixative. Subsequently, grids were blocked with 100 µL PBS containing 1% BSA for 5 min and incubated with ComGC_95-108_ antibodies (provided by Birgitta Henriques-Normark) diluted 1:100 in PBS containing 1% BSA for 1 h [35]. Grids were washed six times with PBS and incubated with protein A conjugated to 10-nm gold particles (Cell Microscopy Center, University Medical Center Utrecht) diluted 1:50 in PBS containing 1% BSA for 45 min. After six washes with PBS, grids were subjected to a final fixation on a 10 µL drop of 1% glutaraldehyde for 5 min, followed by three washes with distilled water. Labeled grids were negatively stained with 1% uranyl acetate aqueous solution for 5 sec. Incubations with glycine, blocking buffer, and antibodies were performed under covered conditions to minimize evaporation.

Specimens were examined using a JEM-1400Flash transmission electron microscope (JEOL) operated at 120 kV. Digital images were captured with a 4k x 4k Rio16 camera (Gatan) and analyzed using the Fiji distribution of ImageJ software (Version 2.16.0/1.54p). Com pilus lengths were measured using a custom macro in Fiji that enhances image contrast and measures the length of freehand lines tracing the Com pilus. For imaging of capsule, sample preparation and imaging were conducted following established protocols [38].

### Dextran exclusion assay

Sample preparation, imaging, and calculations of dextran exclusion width were performed following a previously described protocol [38]. For quantification of capsule thickness, bacterial cells were stained with FITC-dextran with a molecular weight of 2,000 kDa (Sigma-Aldrich). For the capsule porosity assay, various sizes including 4, 20, 40, 60-76 (assumed as 60 for data interpretation), and 150 kDa of FITC-dextran (Sigma-Aldrich) were used instead.

### Statistical analysis

All statistical analyses were performed using GraphPad Prism v10.4.1 (GraphPad Software Inc.). A Shapiro-Wilk test for normal distribution was performed for datasets and specific statistical tests are noted in figure legends.

## ACKNOWLEDGEMENTS

We thank Dr. Birgitta Henriques-Normark for providing ComGC antibodies and Dr. Surya Aggarwal for providing primers SDA35/SDA43. We acknowledge NYU Langone Health Microscopy Lab Alice Liang for consultation and support with electron microscopy.

This work was supported by the NIH (5R01AI150893 and 5R37AI038446 to J.N.W.). The microscopy core is partially supported by NYU Cancer Center Support Grant NIH/NCI P30CA016087.

---

**Fig S1.**
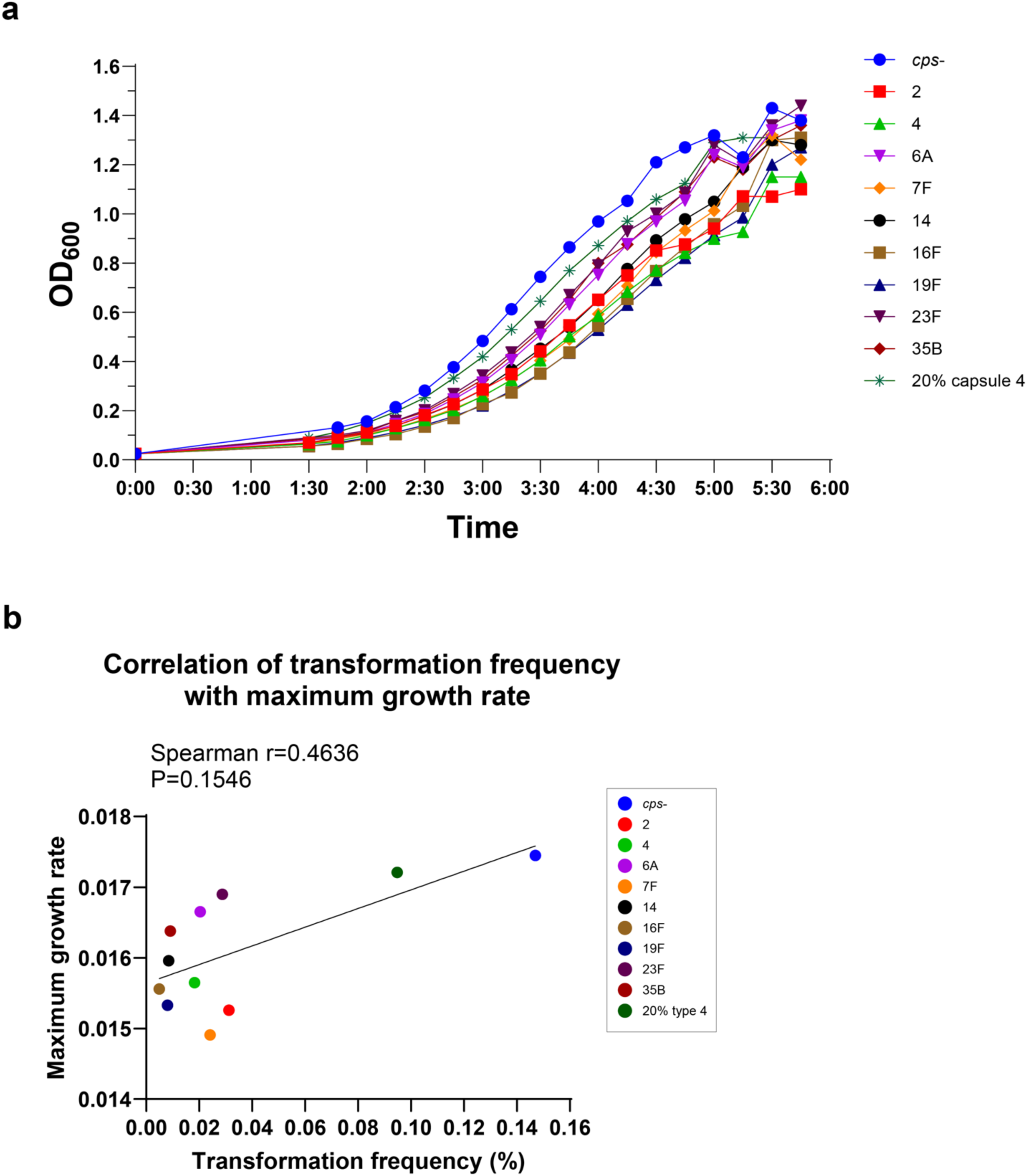
Growth rate of capsule-switch strains. (a) Growth curve of capsule-switch strains. Each strain was cultured in THY (pH=6.6) at 37°C until stationary phase. Optical density at 600 nm (OD_600_) was measured every 15 min. (b) Correlation of transformation frequency with PCR product and maximum growth rate. The maximum growth rate was calculated as the slope of the log phase growth curve on a log scale. Spearman nonparametric correlation coefficients were computed. Strains are designated by their capsule type or amount for brevity.

**Fig S2.**
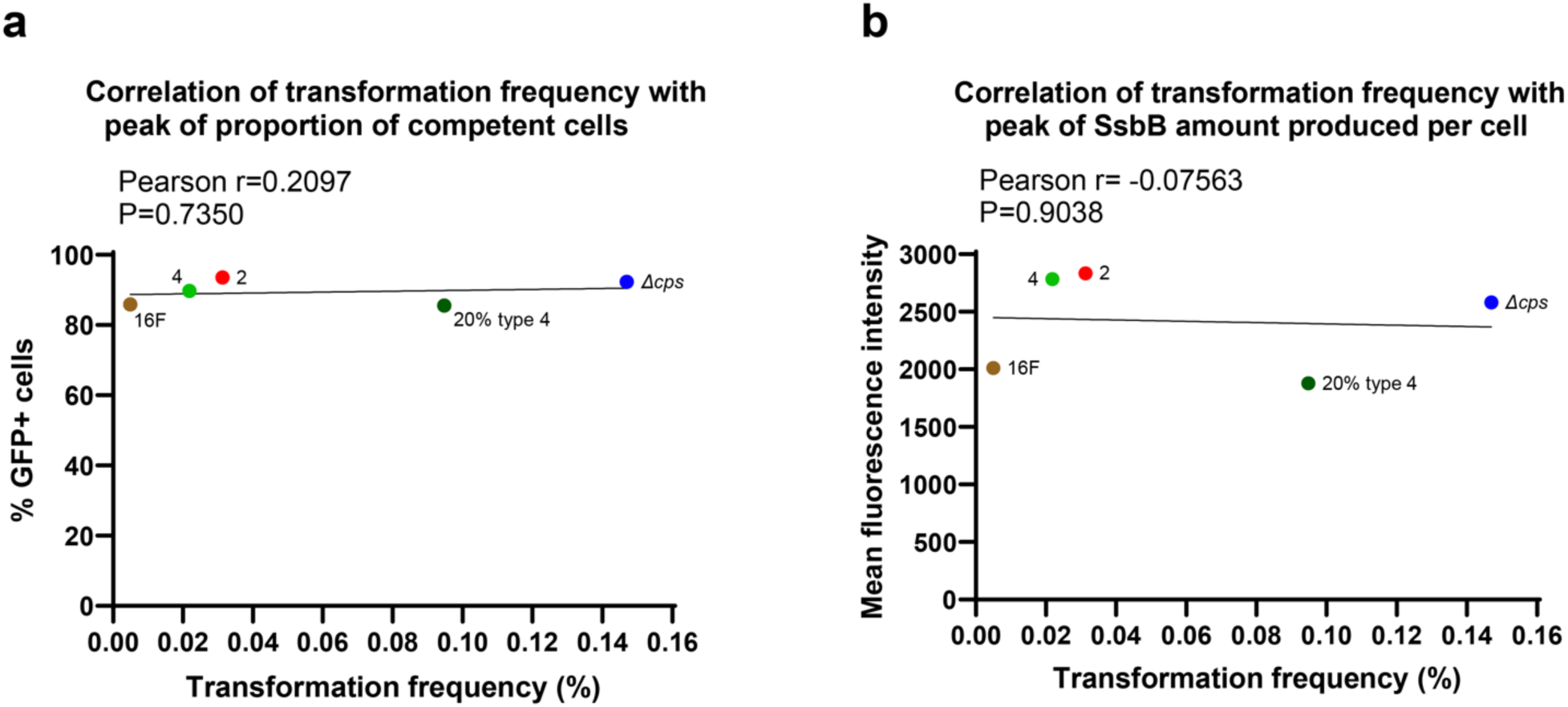
Competence level of five selected representative strains examined by P*_ssbB_-ssbB-gfp* reporters. (a) Correlation of transformation frequency with PCR product and the peak value of proportion of competent cells. Pearson correlation coefficients were computed. (b) Correlation of transformation frequency with PCR product and the peak value of amount of SsbB produced per cell. Pearson correlation coefficients were computed. Strains are designated by their capsule type or amount for brevity.

**Fig S3.**
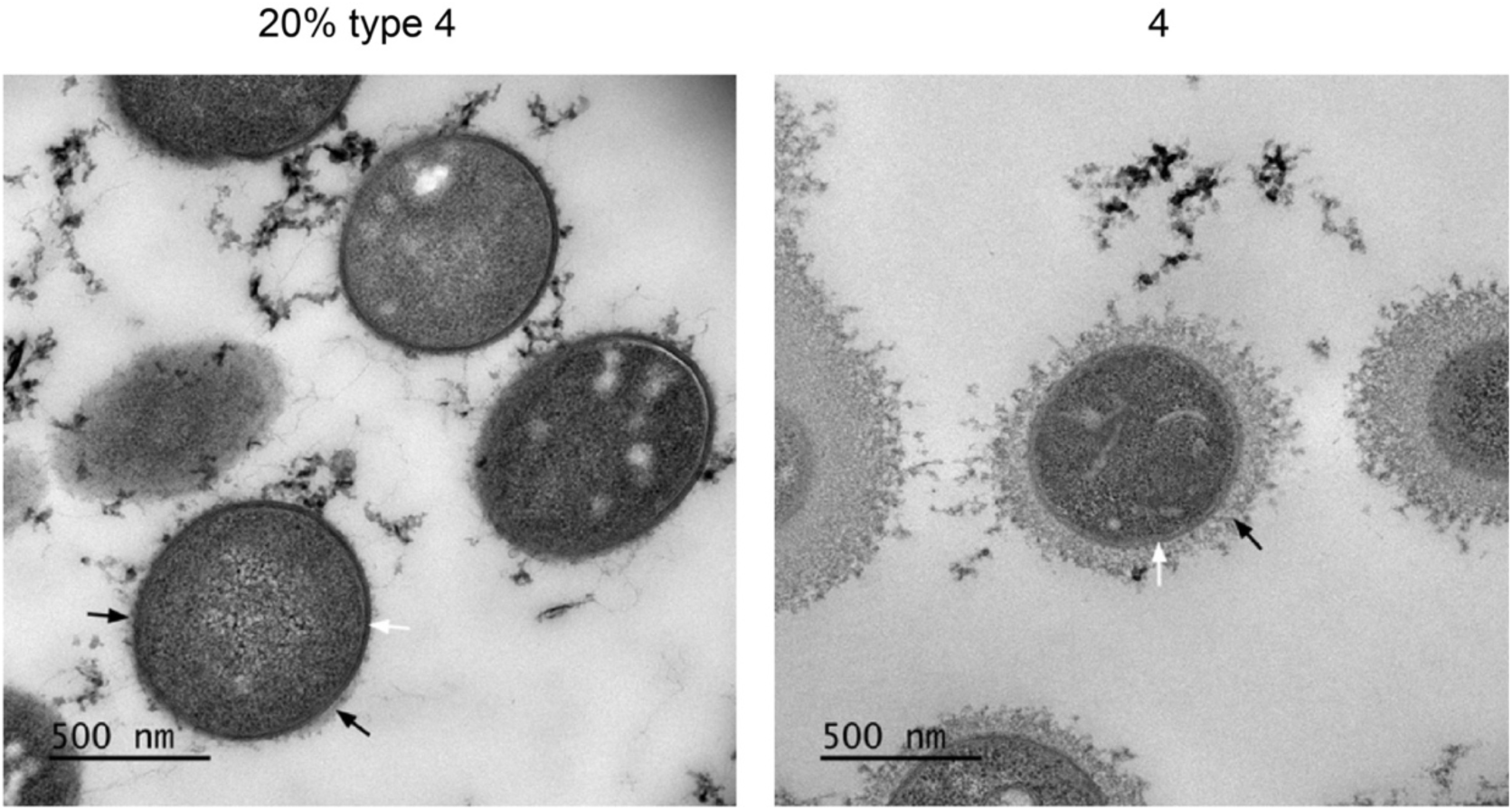
Transmission electron microscopy to visualize the capsule. Bacteria were chemically fixed with lysine acetate added, stained with 0.075% ruthenium red, and embedded in LR White. Black arrows indicate the capsule, and white arrows indicate the cell wall. Scale bar: 500 nm. Strains are designated by their capsule type or amount for brevity.

**Fig S4.**
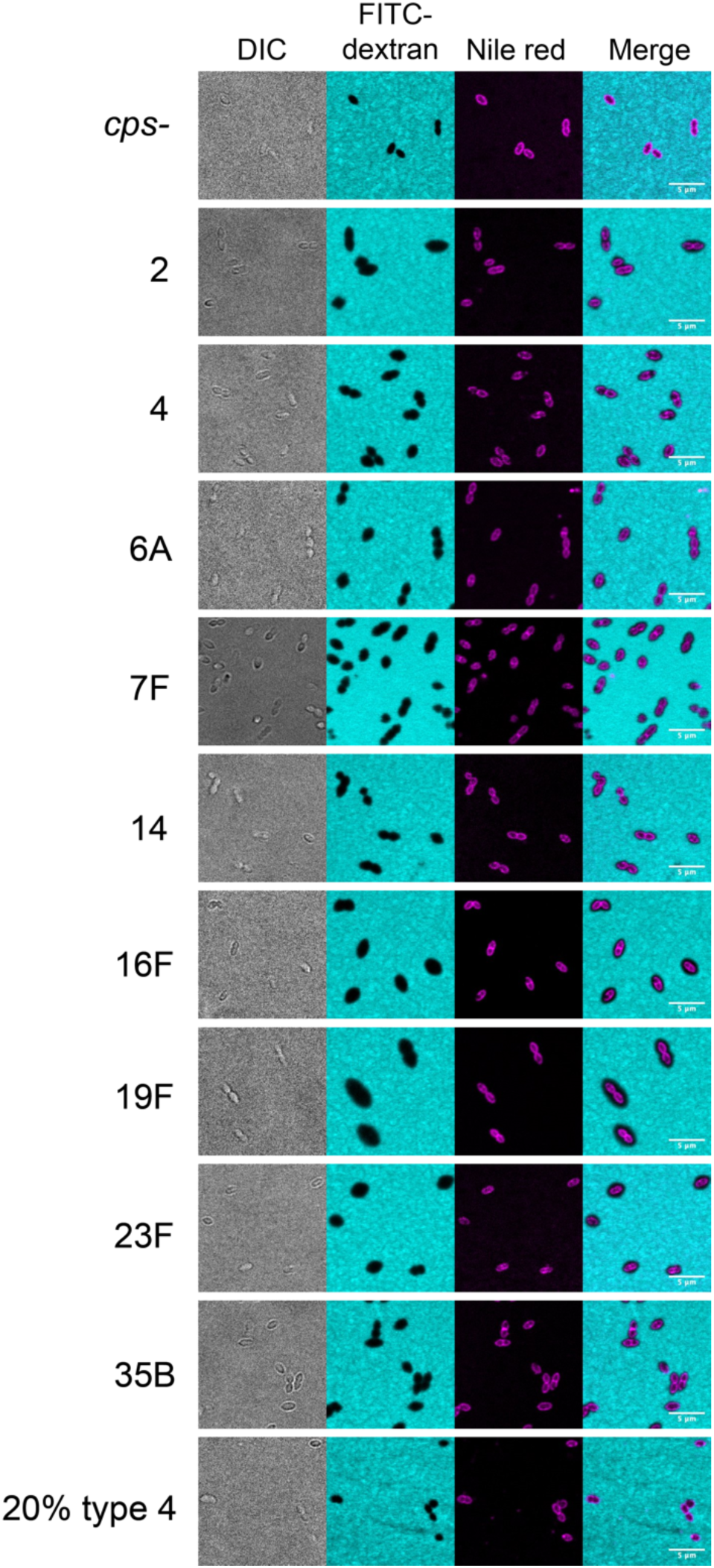
Capsule visualization via fluorescence light microscopy. Bacterial cell body including capsule was visualized as a shadow excluded from the background staining of FITC-conjugated dextran. The cell surface was visualized by staining membrane lipids with Nile red. Strains are designated by their capsule type or amount for brevity.

**Fig S5.**
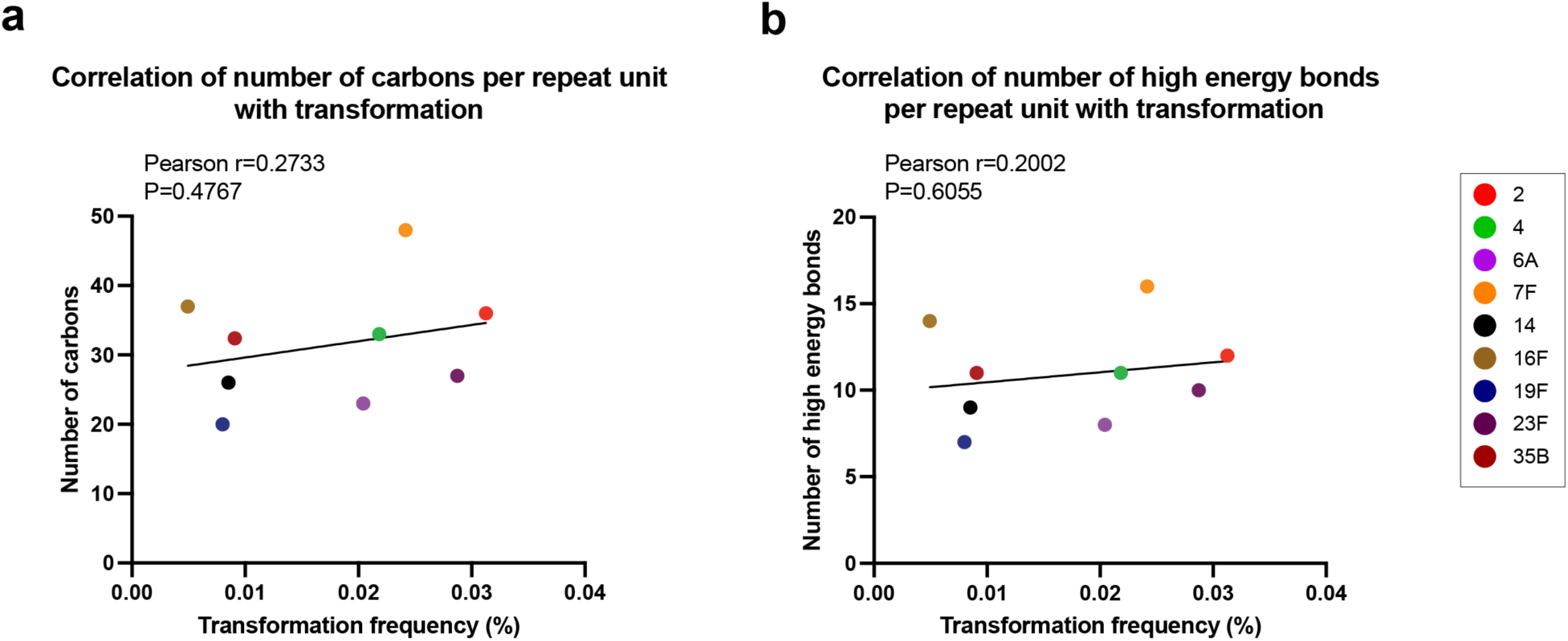
Correlation of transformation frequency with compositional metrics of capsular polysaccharides. (a) Correlation of transformation frequency with number of carbons per polysaccharide repeat unit. Pearson correlation coefficients were computed. (b) Correlation of transformation frequency with number of high energy bonds required to generate one polysaccharide repeat unit. Pearson correlation coefficients were computed. Compositional metrics data were obtained from the previous study [39]. Strains are designated by their capsule type for brevity.

